# ECLIPSER: identifying causal cell types and genes for complex traits through single cell enrichment of e/sQTL-mapped genes in GWAS loci

**DOI:** 10.1101/2021.11.24.469720

**Authors:** John M. Rouhana, Jiali Wang, Gokcen Eraslan, Shankara Anand, Andrew R. Hamel, Brian Cole, Aviv Regev, François Aguet, Kristin G. Ardlie, Ayellet V. Segrè

**Affiliations:** Ocular Genomics Institute and Department of Ophthalmology, Massachusetts Eye and Ear, Boston, MA 02114, USA; Department of Ophthalmology, Harvard Medical School, Boston, MA 02114, USA; Medical and Population Genetics Program, Broad Institute of MIT and Harvard, Cambridge, MA 02142, USA; Klarman Cell Observatory, Broad Institute of MIT and Harvard, Cambridge, MA, 02142, USA; The Broad Institute of MIT and Harvard, Cambridge, MA, USA; Genentech, 1 DNA Way, South San Francisco, CA, USA

## Abstract

**Summary:** ECLIPSER was developed to identify pathogenic cell types and cell type-specific genes that may affect complex disease susceptibility and trait variation by integrating single cell data with known GWAS loci. ECLIPSER maps genes to GWAS loci for a given complex trait based on expression and splicing quantitative trait loci (e/sQTLs) and other functional data, and tests whether the mapped genes are enriched for cell type-specific expression in particular cell types using single-cell/nucleus RNA-seq data from one or more tissues of interest. A Bayesian Fisher’s exact test is used to compute fold-enrichment significance. We demonstrate the application of ECLIPSER on various skin diseases and traits using snRNA-seq of healthy human skin samples.

**Availability and Implementation:** The source code and documentation for ECLIPSER and a Jupyter notebook for generating output tables and figures are available at https://github.com/segrelabgenomics/ECLIPSER. The source code for GWASvar2gene that maps genes to GWAS loci based on e/sQTLs is available at https://github.com/segrelabgenomics/GWASvar2gene. The analysis presented here used data from GTEx (https://gtexportal.org/home/datasets) and Open Targets Genetics (https://genetics-docs.opentargets.org/data-access/graphql-api), but can also be applied to other GWAS variant lists and QTL studies. Data used to reproduce the results of the paper are available in Supplementary data.

## 1 INTRODUCTION

Genome-wide association studies (GWAS) have led to the discovery of tens of thousands of common DNA variant association with a range of complex diseases and traits. However, identifying the underlying causal genes, and the pathogenic cell types through which the genes exert their causal effects on disease remains a major challenge. This is because most associated variants lie in noncoding regions, and due to linkage disequilibrium, are often tagging large genomic regions with multiple variants and genes (Cano-Gamez and Trynka, 2020; Gamazon et al., 2018; Visscher et al., 2017). The Genotype-Tissue Expression (GTEx) Project has been instrumental in identifying variants, including noncoding ones, with regulatory effects associated with gene expression (eQTL) or alternative splicing (sQTLs) across 49 healthy human tissues (GTEx Consortium, 2020; GTEx Consortium, 2017). Expression and splicing QTLs have been shown to significantly contribute to complex disease susceptibility and can be used to prioritize disease-causing genes and pathways (Gamazon et al., 2018; GTEx Consortium, 2020; Barbeira et al., 2021; Finucane et al., 2018). Various strategies have been used to prioritize causal genes in GWAS loci using e/sQTLs, including colocalization analysis (Hormozdiari et al., 2016; Wen et al., 2017), mendelian randomization (Zhu et al., 2016, 2021), and transcriptome-wide association analysis (Gamazon et al., 2015; Gusev et al., 2016; Barbeira et al., 2018), as well as methods that integrate additional functional evidence (Ghoussaini et al., 2021; Weeks et al., 2020). Other methodologies have been developed to propose disease-causing cell types based on genome-wide variant association summary statistics and epigenetic or single cell expression data, such as LD score regression (Finucane et al., 2015; Jagadeesh et al., 2021), enrichment analysis (Slowikowski et al., 2014), and regression models (Calderon et al., 2017; Watanabe et al., 2019). However, to our knowledge, e/sQTL gene mapping and single cell expression data have not yet been combined with known GWAS loci to propose pathogenic cell types and identify genes contributing to disease risk in specific cell type contexts.

Here we developed a method called ECLIPSER (**E**nrichment of **C**ausal **L**oci and **I**dentification of **P**athogenic cells in **S**ingle Cell **E**xpression and **R**egulation data) that maps genes to GWAS loci for a given trait using s/eQTL data and other functional information, and subsequently tests whether the expression of genes mapped to these loci is enriched in specific cell types in a given tissue, when compared to the expression of genes in a background set of GWAS loci of unrelated traits. In addition to proposing key disease-or trait-causing cell types, ECLIPSER prioritizes causal genes in GWAS loci driving the enrichment signal in the specific cell types for experimental follow-up. ECLIPSER is a computational framework that can be applied to single cell or single nucleus (sc/sn)RNA-seq data from multiple tissues and to multiple complex diseases and traits with discovered GWAS associations, and does not require genotype data from the e/sQTL or GWAS studies.

## 2 METHODS

ECLIPSER prioritizes causal cell types and cell type-specific genes and regulatory effects that may influence complex diseases or traits using the following steps: **(1) Defining GWAS variants for complex traits:** Genome-wide significant variant associations (P ≤ 5E-08) for traits of interest are taken by default from Open Targets Genetics (Ghoussaini et al., 2021), which contains tens of thousands of associations for a range of complex traits primarily sourced from the NHGRI-EBI GWAS catalog and the UK Biobank GWAS studies (Supplementary Table 1). For each tissue, a background set of GWAS variants (null set) is compiled that includes the genome-wide significant associations for all traits in Open Targets, excluding the variants associated with the traits of interest and traits relevant to the tested tissue. **(2) Mapping genes to GWAS loci:** Genes whose *cis*-eQTLs or *cis*-sQTLs from 49 GTEx tissues (GTEx Consortium, 2020) are in linkage disequilibrium (LD) with a GWAS variant based on a population-matched reference panel, are mapped to each GWAS variant for both the traits of interest and background traits using GWASvar2gene (see Supplementary Note section B for details). Genes can be further assigned to GWAS loci using ‘bestLocus2Genes’ mapping from Open Targets (Supplementary Table 1), which is based on additional omics data (e.g., pQTLs, Hi-C) and predicted deleterious protein coding variants (Ghoussaini et al., 2021); **(3) LD clumping of GWAS variants:** GWAS variants that are in LD with each other or that share mapped genes are collapsed into a single locus for each trait of interest and for each tissue-specific null set separately, to avoid inflation of enrichment due to LD between GWAS variants (Supplementary Note section B); **(4) Cell type-specificity scoring of GWAS loci:** For each trait of interest, tissue, and cell type combination, a cell type specificity score is computed for each GWAS locus as the fraction of cell type-specific genes mapped to the locus (log2 fold-change > 0.5 and false discovery rate (FDR) < 0.1 used here). Cell type specificity is computed for each gene using differential gene expression analysis, comparing each cell type to all other cells in each tissue (Supplementary Note section C). Similar cell type specificity scores are computed for the null GWAS loci. **(5) Cell type-specific enrichment per GWAS locus set:** Cell type specificity fold-enrichment and significance (p-value) are estimated per GWAS locus set for each trait of interest, tissue, and cell type compared to the tissue-dependent background GWAS loci, using a Bayesian Fisher’s exact test and the 95th percentile of the null GWAS locus scores as the cell type specificity cutoff and uninformative priors (See supplementary Note section D for details). The Bayesian approach enables estimating fold-enrichment confidence intervals for traits with small or zero counts of loci in the contingency table. **(6) Identifying cell type-specific causal genes:** In the significantly enriched cell types (false discovery rate < 0.05), cell type-specific genes mapped to GWAS loci that score above the 95th percentile cutoff and are thus driving the enrichment signal (“leading edge GWAS loci”), are prioritized as causal genes (“leading edge causal genes”) in the particular cell type for the given trait. In cases where a gene is mapped based on e/sQTLs, the regulatory effect is recorded and proposed as a potential underlying causal mechanism (Supplementary Note section E).

### Evaluation of the cell type-specific GWAS enrichment method

We evaluated the performance of the Bayesian Fisher’s exact test in assessing cell type-specific GWAS loci enrichment compared to a permutation-based approach (details in Supplementary Note Section D), using snRNA-seq of 8 GTEx tissues and 21 relevant complex traits (Eraslan et al., 2021) (Supplementary Table 2). We found high concordance of the cell type specificity fold-enrichment (Spearman’s rank correlation coefficient =0.89-0.95) and enrichment p-values (Spearman’s =0.91-0.98) between the two approaches (Supplementary Table 3 and Supplementary Fig. 1).

We further used the null GWAS loci and skin snRNA-seq from GTEx and a second study analyzed below (Reynolds et al., 2021) to evaluate the type 1 error rate of the Bayesian Fisher’s exact test (Supplementary Note section D). We randomly sampled 1,000 sets of 100 GWAS loci each from the null GWAS locus set for skin, and computed cell type-specific enrichment per randomly sampled GWAS locus set, for each of the cell types. We found that on average the fraction of significant enrichment at p-value < 0.01 and p < 0.05 matched the corresponding cutoffs among the various cell types presented in skin (Supplementary Tables 4 and 5).

## 3 IMPLEMENTATION AND PERFORMANCE

ECLIPSER was written in Python. It requires 60Gb of memory to run the LD clumping step and 1Gb of memory for running the enrichment analysis. The estimated run time for generating clumping files for the GWAS loci of a given trait and the null set of GWAS loci is 30 minutes, and the estimated run time for the enrichment analysis of a given trait in a single tissue with 12-16 cell types is 1-3 minutes on a node of 64-bit CPU with 16 cores and 128 GB of RAM.

## 4 EXAMPLE

We applied ECLIPSER to six skin diseases and traits (Supplementary Table 6) using snRNA-seq data from healthy skin samples with 40 defined cell type clusters (Reynolds et al., 2021) (Fig. 1 and Supplementary Note section F). Genes mapped to GWAS loci associated with melanoma, squamous cell carcinoma, and skin pigmentation traits were significantly enriched (Experiment-wide FDR < 0.05) in melanocytes (fold-enrichment (FE)=4.9, 7.2, and 14.1, respectively), while genes mapped to basal cell carcinoma loci were most strongly enriched in macrophages, dendritic cells, and differentiated keratinocytes (FE=2.5-3.3) (Fig. 1, Supplementary Fig. 2A,B and Supplementary Table 8). The inflammatory diseases - atopic dermatitis and psoriasis showed enrichment in both distinct and overlapping immune cell types, with genes associated with atopic dermatitis most strongly enriched in innate lymphoid cells and monocyte-derived dendritic cells (FE=2.4-3.6), and genes associated with psoriasis most strongly enriched in cytotoxic T cells (FE=3.2) (Supplementary Fig. 2C,D and Supplementary Table 8). In addition to proposing disease-critical cell types, this analysis proposes genes and regulatory effects that may contribute to disease susceptibility in specific cell types, as shown for melanoma and melanocytes in Fig. 1B,C, basal cell carcinoma and macrophages in Fig. 1D,E, and other skin traits and cell types in Supplementary Figure 3 and Supplementary Table 8. The application of ECLIPSER to additional complex diseases and traits and 8 snRNA-seq GTEx tissues can be found in (Eraslan et al., 2021).

**Figure 1.**
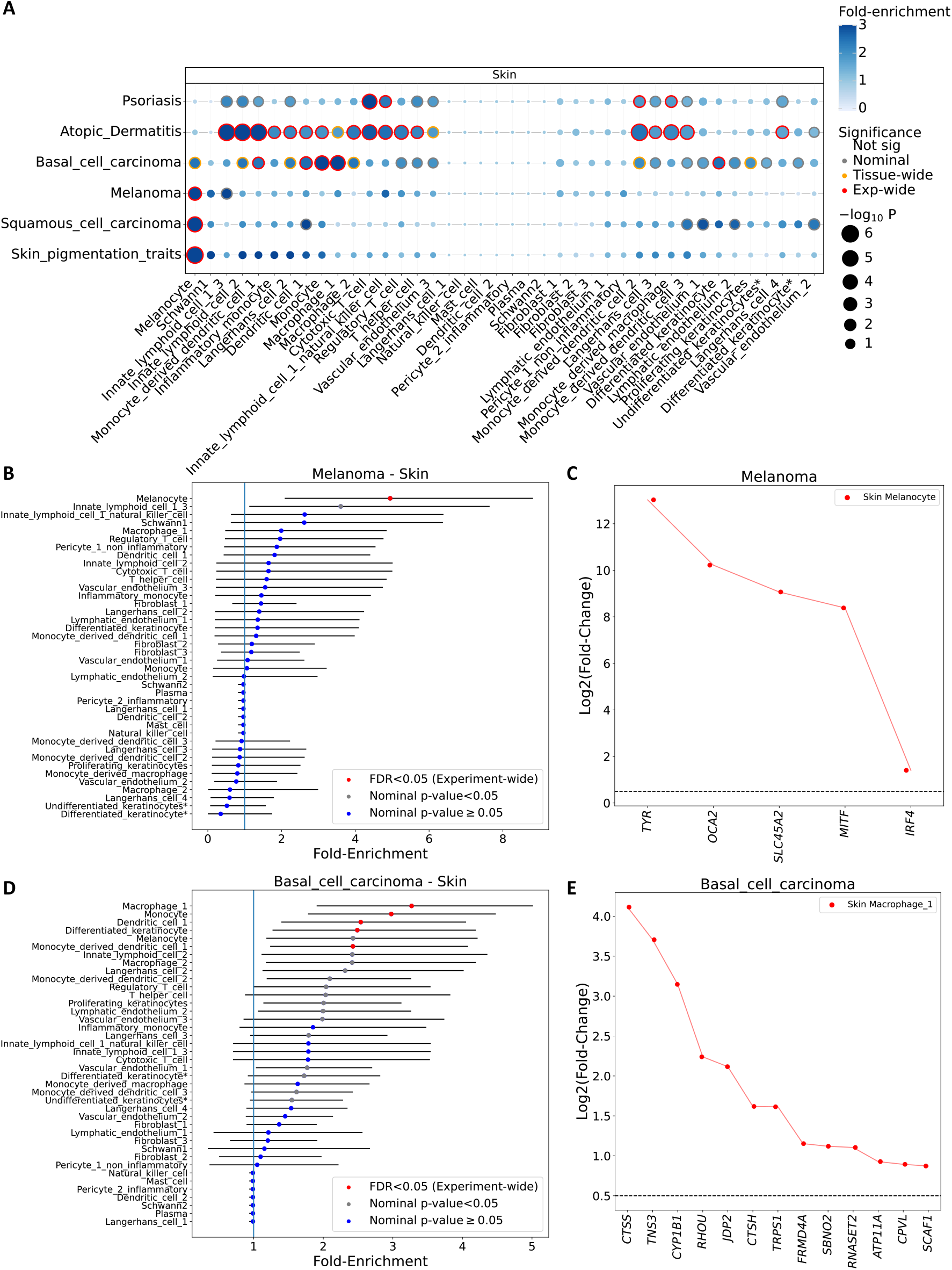
Cell type-specific enrichment of e/sQTL-mapped genes to GWAS loci of skin diseases and traits using skin snRNA-seq expression. (A) Significance (circle size, -log10(P-value)) and fold-enrichment (circle color) of the cell type specificity of GWAS locus sets (rows) of six skin diseases and traits are shown for each of the 40 cell type clusters found in the skin snRNA-seq study (Reynolds *et al*., 2021). Traits (rows) and skin cell types (columns) were clustered based on hierarchical clustering of the euclidean distance between the GWAS locus set cell type-specificity enrichment scores. (B,D) Cell type specificity fold-enrichment (x-axis) in the different skin cell types ranked in descending order are shown for melanoma (B) and basal cell carcinoma (D) GWAS locus sets. Error bars: 95% confidence intervals. Red: experiment-wide significant (Benjamini Hochberg (HB) FDR < 0.05); Gray: nominal significant (P < 0.05); Blue: non-significant (P ≥ 0.05). (C,E) Differential expression (log2(Fold-change), y axis) in the most strongly enriched cell type compared to all other skin cell types for the set of genes (x axis) driving the enrichment signal of melanoma GWAS loci in melanocytes (C) and basal cell carcinoma GWAS loci in macrophages (Macro_1) (E).

## ACKNOWLEDGEMENTS

This work was supported by the Chan Zuckerberg Initiative (CZI) Seed Network for the Human Cell Atlas awards CZF2019-002459 (A.V.S, J.M.R) and 2019-02455 (K.G.A.), National Institutes of Health, National Eye Institute R01 EY031424-01 (A.V.S, J.M.R, A.R.H., B.C.), and NIH, NHGRI 5U41HG009494 (S.A., F.A., K.G.A.), and funds from the Manton Foundation, Klarman Family Foundation and HHMI (A.R.).

