## Supplementary Notes and Figures for "ECLIPSER: identifying causal cell types and genes for complex traits through single cell enrichment of e/sQTL-mapped genes in GWAS loci"

November 24, 2021

### Supplementary Notes

Here we provide more details on the ECLIPSER methodology and parameters used in the applications of ECLIPSER presented in this paper.

#### A. Defining GWAS variants for traits of interest and background traits

In this study GWAS variants were compiled from Open Targets Genetics (Ghoussaini *et al.*, 2021), which contains associations from the NHGRI-EBI GWAS catalog and UK Biobank GWAS, using their API (details in (Eraslan *et al.*, 2021)). GWAS variants for traits of interest can also be taken from any available GWAS database or study. Only genome-wide significant variants (P < 5E-08) were considered. For each trait of interest, all GWAS that corresponded to the given trait were considered together. For each tissue, a background set of GWAS variants (null set) was assembled from the genome-wide significant associations of all traits in Open Targets Genetics (Supplementary Table 1), excluding variants associated with traits whose pathophysiology is believed to be relevant to the tested tissue, which were manually compiled. ECLIPSER provides the option to restrict to a particular ancestral background. In this work, we restricted to GWAS that contained samples of European ancestry.

#### B. Mapping genes to GWAS variants and LD clumping

In the analyses in this paper, genes were mapped to trait-associated loci for the selected and background (null) traits, using a 95% credible set of fine-mapped *cis*-eQTLs and *cis*-sQTLs from each of the 49 GTEx tissues (v8) computed using DAP-G and the European subset of GTEx samples (GTEx Consortium, 2020; Barbeira *et al.*, 2021). e/sQTLs from all 49 tissues were used for the mapping of genes to GWAS loci for all traits, not only from the relevant tissues per trait, as a large fraction of e/sQTLs have been shown to be shared across multiple tissues and the QTL discovery power varies substantially between tissues due to differences in tissue sample sizes ( 70-700 samples/tissue; GTEx Consortium, 2020). The e/sQTL-based gene mapping was performed using the GWASvar2gene tool (`https://github.com/segrelabgenomics/GWASvar2gene`) based on the following steps: (i) Variants in linkage disequilibrium (LD) ($r2 \geq 0.8$) with each of the GWAS variants were identified using the European subset of the GTEx whole genome sequencing (WGS) variant calls as the reference panel (GTEx Consortium, 2020; Barbeira *et al.*, 2021) and PLINK 2.0 (*Plink –bfile Reference_panel_chr_files –r2 –ld-snp-list variant_list_file –ld-window-kb 5000 –ld-window-r2 0.8 –ld-window 99999*). (ii) If a GWAS variant was not present in the GTEx WGS variant call set, LD proxy variants to the GWAS variant at $r2 > 0.8$ were searched in the 1000 Genomes Project reference panel (Zheng-Bradley *et al.*, 2017), using the European samples. If found, the proxy variants were subsequently checked in the GTEx panel. (iii) GWAS associations whose variants or LD proxy variants were significant eVariants or sVariants in any of the 49 GTEx tissues were assigned the corresponding eGene(s) and/or sGene(s) to their locus. (iv) Genes were further assigned to GWAS variants based on the 'bestLocus2Genes' mapping in Open Targets Genetics (Supplementary Table 1), which includes additional omics data (e.g., protein QTLs, Hi-C) and predicted deleterious protein coding variants that are in LD with the respective GWAS variant (Ghoussaini *et al.*, 2021). A given GWAS variant may have more than one gene mapped to it. (v) Two or more GWAS variants with one or more common LD proxy variants or with shared mapped genes were collapsed into a single GWAS locus to avoid inflation of GWAS locus-set enrichment due to multiple GWAS variants tagging the same causal haplotype. The LD clumping was performed separately for the GWAS variants associated with each trait of interest, and all GWAS variants for the tissue-specific background (null) set of loci.

#### C. Cell type-specific gene expression analysis

Cell type-specific gene expression analysis was performed for all gene by cell type combinations, comparing the expression of each gene in each cell type to that across all other cells. Different methods that can be used to perform single cell differential gene expression analysis, such as Wilcoxon rank sum test, Welch's t-test, or regression-based models (Finak *et al.*, 2015). Here we used the Welch's t-test, after correcting for covariates using ComBat (Leek *et al.*, 2012) (see section F below) for the skin snRNA-seq study (Reynolds *et al.*, 2021), as previously applied to snRNA-seq of 8 GTEx tissues in (Eraslan *et al.*, 2021). Cell type-specific gene expression was defined in this study as log2 fold-change $> 0.5$ and false discovery rate (FDR) $< 0.1$. The tool has the option of filtering out genes per cell type if they are not expressed in a minimum percentage of cells of the given cell type.

#### D. GWAS cell type enrichment analysis

In this section, we restrict to one trait of interest, without introducing the notation for trait. We first defined a GWAS locus cell type specificity statistic. Let $i = 1, ..., n$ be the index of a GWAS locus, $j = 1, ..., n_i$ be the index of a gene mapped to GWAS locus $i$, and $c = 1, ..., p$ be the cell type index. Let $x_{jc}$ be the differential gene expression test statistic, log2(fold change) (log2FC), and $y_{jc}$ be the false discovery rate (FDR) from the cell type specific differential gene expression analysis, for gene $j$ and cell type $c$.

For GWAS locus $i$ and cell type $c$, the cell type specificity statistics is defined as the fraction of genes mapped to the locus that are cell type specific:

$$s_{ic} = \frac{\sum_{j=1}^{n_i} \mathbb{1}_{(x_{jc} \geq T_x, y_{jc} \leq T_y)}}{n_i}, \tag{1}$$

where $T_x$ and $T_y$ are the thresholds for log2FC and FDR, respectively, and $\mathbb{1}$ is the indicator variable.

Next, a GWAS locus set test statistic is computed for cell type $c$, as the average of GWAS locus cell type specificity statistics of all loci that belong to the given trait, as follows:

$$s_c = \frac{\sum_{i=1}^{n} s_{ic}}{n} \tag{2}$$

Finally, to test whether the expression of genes in GWAS loci associated with a trait is enriched in specific cell types more than expected by chance, we performed Fisher's exact test to compute a fold-enrichment and assess enrichment significance. A GWAS locus was considered to have high cell type specificity if its specificity score in equation (1) was equal to or higher than the 95th

| | Number of loci $\geq$ 95 percentile of specificity statistics from null set | Number of loci $<$ 95 percentile of specificity statistics from null set | Total |
|---|---|---|---|
| GWAS locus set | c | d | $n_1$ |
| Null set | a | b | $n_2$ |

percentile of the locus cell type specificity statistics of the null set of loci. We constructed the following contingency table:
where $a$ and $b$ are the numbers of GWAS loci $\geq$ 95th percentile and $<$ 95th percentile of the cell type specificity statistics from the null loci, respectively, for the null set of GWAS loci. $c$ and $d$ are the corresponding numbers for the GWAS locus set of the trait of interest. $n_1$ and $n_2$ are the total number of GWAS loci for the trait of interest and null set, respectively. The *fold-enrichment* estimate can be defined as the ratio of the fraction of cell type-specific loci in the GWAS locus set and the fraction of cell type-specific loci in the null set.

$$\text{fold enrichment} = \frac{c/n_1}{a/n_2}. \tag{3}$$

**We tested two approaches to estimating the enrichment significance:**

##### Permutation-based approach

The same number of loci as the number of GWAS loci of each selected trait are randomly sampled with replacement from the background loci $N$ times to generate an empirical null distribution of loci sets. For each cell type and permutation, the number of loci with a cell type specificity score $\geq$ 95th percentile of all null set scores is computed, and compared against the value $a$ of the trait of interest. Since the number of loci of the trait of interest and the null set are the same, the fold-enrichment from one permutation is $\frac{c}{a}$. The 95% confidence interval (CI) can be computed from the $N$ permutations as the 2.5th and 97.5th percentiles of all permutations' fold-enrichment values. The empirical p-value of the fold-enrichment is the fraction of permutations whose fold-enrichment is equal to or higher than the trait's observed fold-enrichment.

##### Bayesian-based approach

However, in the case of small numbers of GWAS loci, the counts in the contingency table might be small or even zero, thus making the fold-enrichment confidence intervals unreliable or impossible to construct. Therefore, we adopted a Bayesian approach to address this problem. Specifically, let $\theta_1$ and $\theta_2$ be the fraction of loci in the GWAS locus set and null locus set, respectively, whose cell type specificity statistic is $\geq$ 95 percentile of the null set cell type specificity statistics.

The observed numbers of loci $\geq$ 95 percentile of the statistics from the null, in the GWAS locus set $c$ and in the null locus set $d$ can be modeled as Binomial distributions, respectively:

$$c \sim Bin(n_1, \theta_1), a \sim Bin(n_2, \theta_2) \tag{4}$$

Assuming uniform priors for $\theta_1$ and $\theta_2$:

$$\theta_1 \sim Unif(0,1), \theta_2 \sim Unif(0,1), \tag{5}$$

the posterior distributions of $\theta_1$ and $\theta_2$ are:

$$\theta_1 \sim Beta(c+1, d+1), \theta_2 \sim Beta(a+1, b+1). \tag{6}$$

The fold-enrichment is thus estimated as:

$$\text{fold-enrichment} = \frac{c/n_1}{a/n_2} = \frac{\theta_1}{\theta_2}. \tag{7}$$

Monte Carlo samples of ($\theta_1$, $\theta_2$) can be drawn from the posterior distributions in equation (6) $N$ times to compute the point estimate, 95% CI, and an empirical Bayesian p-value of the fold-enrichment.

To determine enrichment significance, multiple hypothesis correction is applied tissue-wide (correcting for the number of cell types tested per tissue-trait pair) or experiment-wide (correcting for the number of cell type by tissue by trait combinations tested), using the Benjamini-Hochberg false discovery rate (FDR).

##### Comparison of Bayesian and Permutation enrichment methods

To evaluate the performance of the Bayesian Fisher's exact test to that of the permutation-based test, we used snRNA-seq of 8 GTEx tissues and 21 relevant complex traits (Eraslan *et al.*, 2021) (Supplementary Table 2). We applied the permutation approach to all trait by tissue comparisons using $N$=10,000 permutations, and compared the permutation-based fold-enrichment and enrichment p-value to that from the Bayesian approach computed in (Eraslan *et al.*, 2021). We found high concordance of the cell type specificity fold-enrichment (Spearman's rank correlation coefficient = 0.89, $P$<1E-16) and enrichment p-values (Spearman's rho =0.91, $P$<1E-16) between the two approaches when analyzing GWAS with at least 5 GWAS loci (Supplementary Fig. 1A,B and Supplementary Tables 2 and 3) and even higher concordance (Spearman's rho=0.95 and 0.98, $P$<1E-16) for the fold-enrichment and enrichment p-value, respectively, when considering a subset of 17 traits with at least 10 GWAS loci (Supplementary Fig. 1C,D).

##### Estimating type 1 error rate of Bayesian Fisher's exact test

We evaluated the type 1 error of the Bayesian-based enrichment method in ECLIPSER using permutation analysis of null GWAS loci and cell type-specific skin snRNA-seq expression from GTEx (Eraslan *et al.*, 2021) and a second study (Reynolds *et al.*, 2021) analyzed as an example below (section F). Specifically, we randomly selected 100 GWAS loci from the null set of loci, which represent the loci for the 'trait of interest', and ran Bayesian Fisher's exact test on the locus set and each of the cell types, to obtain nominal enrichment p-values for each cell type and permutation. We repeated this process 1,000 times and computed the fraction of permutations whose enrichment p-values passed 0.01 or 0.05 cutoffs. The results per cell type and cutoff are shown in Supplementary Tables 4 and 5 for the 16 cell types in GTEx skin and 40 cell types in Reynolds *et al.* skin, respectively.

We found that for most of the cell types, the fraction of permutations with significant enrichment roughly matched the nominal p-value cutoffs of 0.01 and 0.05, with an average p-value of 0.011 and 0.045, respectively, for GTEx skin (Supplementary Tables 4), and 0.0097 and 0.054, respectively, for Reynolds *et al.* skin (Supplementary Tables 5). While some of the deviations are due to random noise, we observed that in cell types where the fraction of permutations that passed the given p-value cutoff was 0 (e.g., adipocyte, immune T cell, and immune mast cell in GTEx), the 95 percentile of cell type specificity statistics from the null loci was 0, suggesting that these cell types have low numbers of genes that are cell type specific (Supplementary Table 4). The enrichment p-values for all 1,000 randomly sampled null GWAS loci for these cell types were non-significant (p-value and fold-enrichment were very close to 1), as both $b$ and $d$ in the contingency table are 0. For cell types that have low levels of cell type specificity this is inevitable, leading to more conservative results.

#### E. Prioritizing cell type-specific causal genes for complex traits

For significantly enriched cell types for a given trait, ECLIPSER identifies the set of GWAS loci that are driving the enrichment signal ("leading edge GWAS loci" ), defined as the loci with cell type specificity scores above the 95th percentile of scores in the null set of loci. Cell type-specific genes (defined in section C) mapped to these leading edge GWAS loci are proposed to be causal to the given trait in the enriched cell type ("leading edge genes"). This analysis further proposes regulatory mechanisms through which the potentially causal genes may be contributing to disease risk or trait variation, in cases where eQTLs or sQTLs were used to map the genes to the leading edge GWAS loci.

#### F. Example: Application of ECLIPSER to skin-related diseases and traits and skin snRNA-seq dataset

We demonstrate the utility of ECLIPSER on six skin traits: melanoma, non-melanoma skin cancers (basal cell carcinoma and squamous cell carcinoma), atopic dermatitis, psoriasis, and skin pigmentation traits, using snRNA-seq data from healthy skin samples (Reynolds *et al.*, 2021); https://developmentcellatlas.ncl.ac.uk/datasets/hca_skin_portal/). The GWAS that correspond to these six skin diseases and traits in Open Targets Genetics are listed in Supplementary Table 6. For the skin single nucleus gene expression data, we retrieved a final, annotated Scanpy AnnData object from https://zenodo.org/record/4310074.YZbUnC1h1Yi (submission.h5ad). We filtered for patients with an indicated "Healthy" status, and only included cells that had "final_clustering" annotations. We computed log(TP10K+1) using the "sc.pp.normalize_total" and "sc.pp.log1p" functions in Scanpy, and performed Combat (Leek *et al.*, 2012) to correct for "donor_id" using "sc.pp.combat" implemented in Scanpy. Finally, we performed differential gene expression analysis for all genes by 40 cell types defined in this snRNA-seq skin study, using Welchś t-test implemented in the "sc.tl.rank_genes_groups" function of Scanpy with the "t-test_overestim_var" option (results in Supplementary Table 7), as applied to the GTEx tissues (Reynolds *et al.*, 2021). Cell type-specific gene expression was defined based on the following criteria: log2 fold-change $> 0.5$, false discovery rate (FDR) $< 0.1$, and the gene is expressed in at least 5% of cells in the given cell type. The null set of GWAS loci was generated by excluding variants associated with all skin-related traits listed in Supplementary Table 6.

The ECLIPSER results for the different skin diseases and traits are shown in Supplementary Table 8, and displayed in Figure 1 and Supplementary Figures 2 and 3. Significant enrichment was found in previously described and less well established cell types for the different skin traits. Putative causal genes proposed for the different traits and the top enriched cell types are shown in Figure 1C,E and Supplementary Figure 3. Notably, the genes driving the enrichment in melanocytes for melanoma, squamous cell carcinoma, and skin pigmentation traits are almost identical, while only about a third of the genes driving the enrichment signal in cytotoxic T cells in psoriasis and atopic dermatitis are common: *BACH2*, *CD247*, *ETS1*, *FASLG*, *RUNX3*, and *TAGAP* (Supplementary Figure 3).

#### G. Conclusions

The ECLIPSER GWAS cell type enrichment method enables to identify key cell types that may contribute to the pathophysiology of a given complex disease or trait, and prioritize disease or trait-causing genes in GWAS loci that often contain multiple genes, in specific cell type contexts. Functional experiments in relevant cellular contexts will be needed to confirm the hypotheses generated by this method.

By comparing the cell type specificity scores of GWAS loci associated with a given trait to a null distribution of GWAS locus scores, as implemented in ECLIPSER, we can detect cell types that are specific to the trait of interest and not common to all traits. In doing so, we also correct for potential confounding factors that may be shared across GWAS loci. Computing cell type specificity scores between cells within each tissue separately accounts for potential biases within each tissue, and enables to compare cell type enrichment scores across tissues. One limitation of this method is that genes that are ubiquitously expressed across many or all cell types in a given tissue are not considered as potential causal genes in this analysis, though some genes may contribute to disease risk or trait variation through multiple cell types or through one cell type but still be expressed in multiple cell types.

#### H. Software

The source code and detailed documentation on how to run ECLIPSER and a Python Jupyter notebook for generating the output tables and figures is available at `https://github.com/segrelabgenomics/ECLIPSER`. The source code and documentation on how to run GWAS-var2gene, a tool that maps genes to GWAS loci based on e/sQTLs that are in LD with each corresponding GWAS variant is available at `https://github.com/segrelabgenomics/GWASvar2gene`. Both tools were written in python.

#### I. Data

The GTEx eQTLs and sQTLs from release v8 were downloaded from `https://gtexportal.org/home/datasets`. The variant-to-gene ('bestLocus2Genes') mapping from Open Targets Genetics was downloaded from `https://genetics-docs.opentargets.org/data-access/graphql-api` using their API as described in (Eraslan *et al.*, 2021). The snRNA-seq data of the healthy skin samples analyzed in this paper were taken from: `https://zenodo.org/record/4310074#.YZbUnC1h1Y` (Reynolds *et al.*, 2021).

#### J. URLs

**GTEx v8:** https://gtexportal.org/home/datasets)
**Open Targets Genetics:**
https://genetics-docs.opentargets.org/data-access/graphql-api
**GWASvar2gene:** https://github.com/segrelabgenomics/GWASvar2gene
**ECLIPSER:** https://github.com/segrelabgenomics/ECLIPSER
**Reynold *et al.* 2021 skin snNRA-seq dataset:** `https://zenodo.org/record/4310074#.YZbUnC1h1Y`

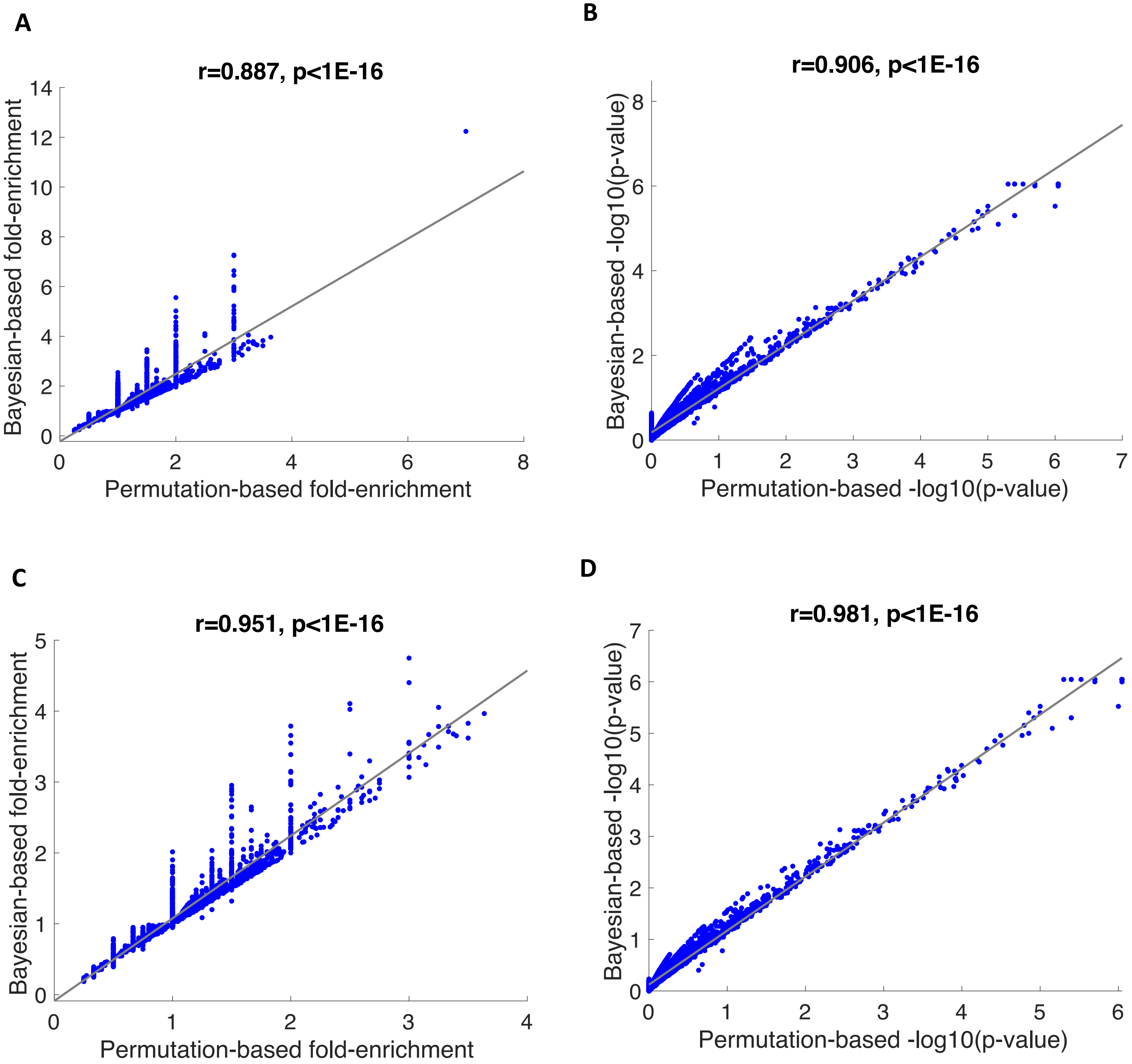


**Supplementary Figure 1. Correlation between Permutation- and Bayesian-based GWAS cell type specificity enrichment statistics.** The correlation of the cell type specificity fold-enrichment (A,C) and enrichment significance, –log10(P-value) (B,D) between the Bayesian and permutation-based Fisher's exact test is shown for 8 snRNA-seq GTEx tissues tested by 21 complex traits with at least 5 GWAS loci (A,B) or a subset of 17 traits with at least 10 GWAS loci (C,D). r, Spearman's rank correlation coefficient.

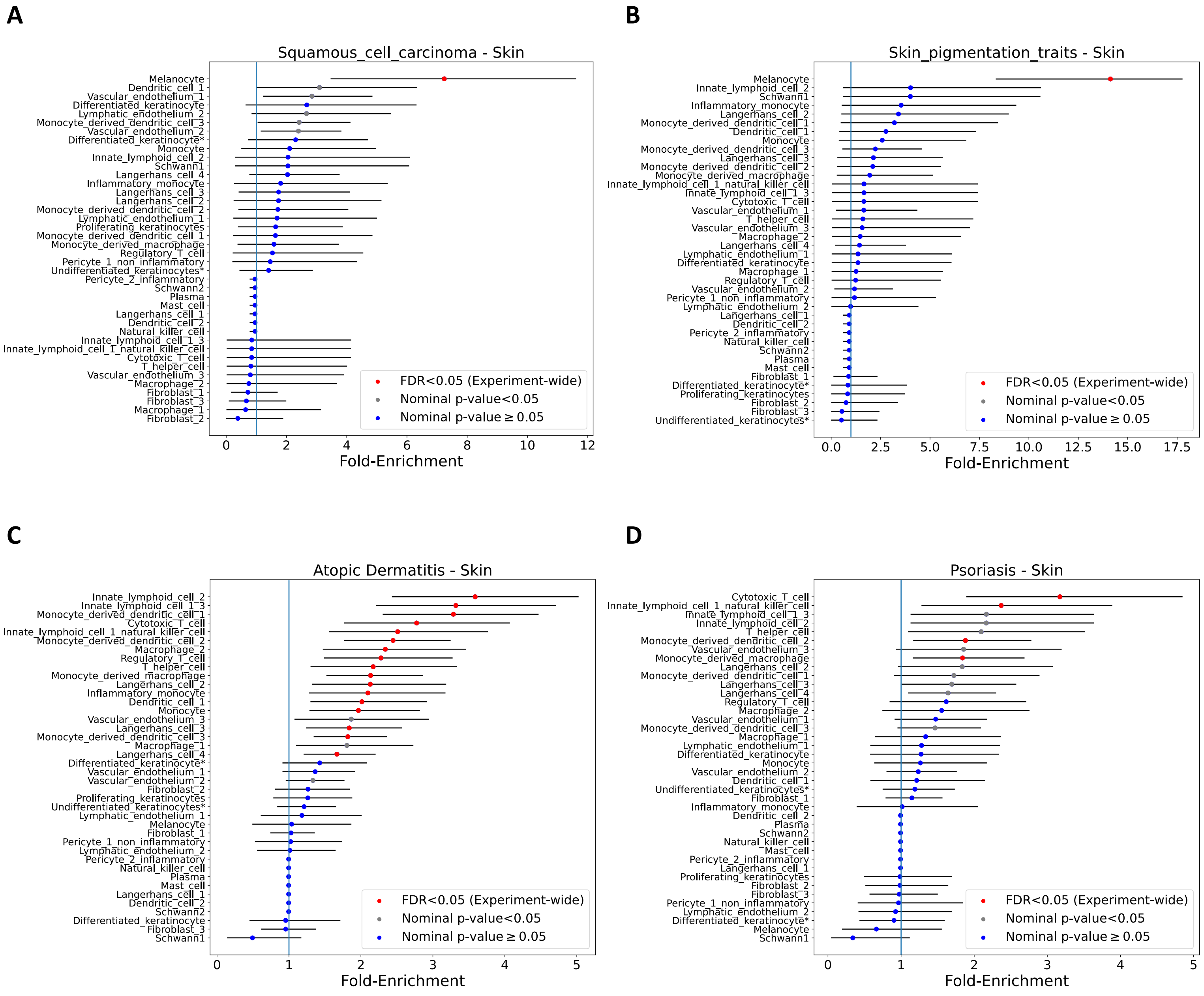


**Supplementary Figure 2. Fold-enrichment of cell type-specific expression in GWAS loci of skin diseases and traits in skin snRNA-seq study**. The top enriched cell types are shown for Squamous cell carcinoma (A), skin pigmentation traits (B), Atopic dermatitis (C), and Psoriasis (D) using snRNA-seq data from healthy skin samples (Reynolds *et al.*, 2021). Error bars represent 95% credible intervals. Red: experiment-wide significant (Benjamini Hochberg (HB) FDF<0.05); Gray: nominal significance (P<0.05).

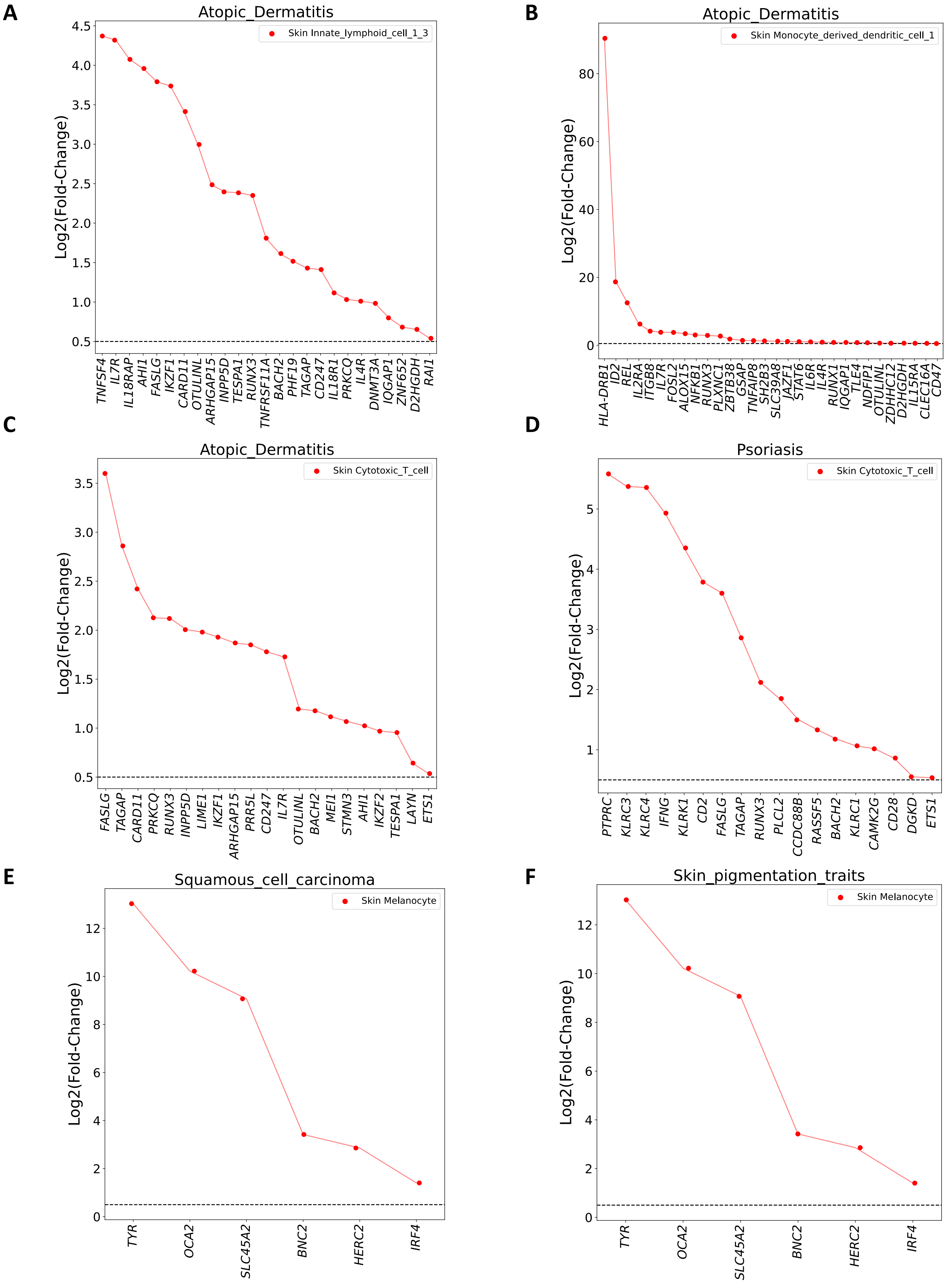


**Supplementary Figure 3. Genes driving the cell type enrichment signal for GWAS loci of skin diseases and traits in skin snRNA-seq.** Differential expression (log2(Fold-change), y axis) in the most strongly enriched skin cell types compared to all other cells for the cell type-specific genes (x axis) driving the enrichment signal for atopic dermatitis (A-C), Psoriasis (D), squamous cell carcinoma (E), and skin pigmentation traits (F) GWAS loci in the corresponding enriched cell type (listed in legend).
